# The central limit theorem for the number of mutations in the genealogy of a sample from a large population

**DOI:** 10.1101/2025.01.23.634620

**Authors:** Yun-Xin Fu

## Abstract

The number *K* of mutations identifiable in a sample of *n* sequences from a large population is one of the most important summary statistics in population genetics and is ubiquitous in the analysis of DNA sequence data. *K* can be expressed as the sum of *n* −1 independent geometric random variables. Consequently, its probability generating function was established long ago, yielding its well-known expectation and variance. However, the statistical properties of *K* is much less understood than those of the number of distinct alleles in a sample. This paper demonstrates that the central limit theorem holds for *K*, implying that *K* follows approximately a normal distribution when a large sample is drawn from a population evolving according to the Wright-Fisher model with a constant effective size, or according to the constant-in-state model, which allows population sizes to vary independently but bounded uniformly across different states of the coalescent process. Additionally, the skewness and kurtosis of *K* are derived, confirming that *K* has asymptotically the same skewness and kurtosis as a normal distribution. Furthermore, skewness converges at speed 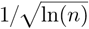 and while kurtosis at speed 1 /ln *n*. Despite the overall convergence speed to normality is relatively slow, the distribution of *K* for a modest sample size is already not too far from normality, therefore the asymptotic normality may be sufficient for certain applications when the sample size is large enough.

---

The number *K* of mutations in the genealogy of a sample of size *n* is one of the most well-known summary statistics in population genetics. Under the infinite-sites model, *K* corresponds to the number of segregating sites (or polymorphic sites) (e.g., Ewens (2004)). Watterson (1975) established the finite-sample probability generating function for *K* and derived its expectation and variance, given by

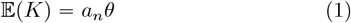

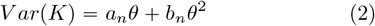

where θ = 4*Nµ* with *N* as the effective population size and *µ* as the mutation rate per sequence per generation, and

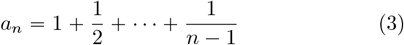

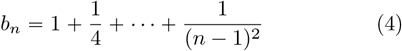

Under coalescent theory (Kingman, 1982a,b; Hudson, 1983; Tajima, 1983), these two moments can be derived more concisely through conditional expectations. The informativeness of *K* and elegance of coalescent theory have stimulated a wide range of methodological/theoretical studies on *K*, including its distribution(Tavaré, 1984), the estimation of *θ* (Fu and Li, 1993b; Pluzhniko and Donnelly, 1997), the inference about the age of the most recent common ancestor of a sample (Fu and Li, 1996; Fu, 1996a; Donnelly et al., 1996; Weiss and von Haeseler, 1996; Tavaré et al., 1997), the partition of *K* into components (Fu and Li, 1993a; Fu, 1995, 2022), and the test of neutrality (Tajima, 1989; Fu and Li, 1993a; Fu, 1996b, 1997; Fay and Wu, 2000; Innan et al., 2005; Zeng et al., 2006; Achaz, 2009; Ronen et al., 2013). Despite these (and many other studies not cited here) significance progress, two aspects of *K* have received little attention. The first is its asymptotic distribution as the sample size *n* → ∞, which has both theoretical and practical implications. The second is its higher moments, particularly skewness and kurtosis, which are valuable for gaining a deeper understanding of its finite-sample properties.

The role of *K* under the infinite-sites model is analogous to that of the number *k* of distinct alleles in a sample under the infinite-alleles model. However, the statistical properties of *k* have been more thoroughly characterized. In deriving the celebrated Ewens sampling formula (Ewens, 1972; Karlin and McGregor, 1972), all moments of *k* were established, leading Ewens to conclude that *k* follows an asymptotic normal distribution. Recognizing that the variance of *K* is approximately equal to its expectation, Watterson (1975) suggested that *K* follows an approximately Poisson (or normal) distribution for large samples. However, this claim has never been rigorously proven. For a distribution to be asymptotically normal, a sufficient condition is that all power moments equal asymptotically to those of a normal distribution, which is one of the earliest versions of the Central Limit Theorem (CLT) (e.g. Theorem 30.2 in Billingsley (1986)). To date, only the first two moments of *K* have been explicitly derived.

Fu and Li (1993a) demonstrated that Watterson’s estimate of *θ*

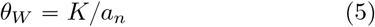

has asymptotically the same variance as a hypothetical Maximum Likelihood Estimator of *θ*, suggesting that *θ*_*W*_ may exhibit asymptotic normality. However, this remains speculative, as the asymptotic distribution of *K* has not yet been formally established. While the asymptotic distribution of *K* may be of more theoretical interest, the higher moments of *K* for finite sample sizes, particularly its skewness and kurtosis, are of practical utility and offer a more complete characterization of *K*.

The purpose of this paper is to prove the asymptotic normality of *K* by applying a specific version of CLT from Probability Theory. Additionally, it derives the skewness and kurtosis of *K* under the assumption that the population evolves according to the Wright-Fisher model, either with a constant effective population size or with varying (independently but uniformed bounded) effective sizes across different states of the sample’s coalescent process.

## The Central Limit Theorem for *K*

Consider a sample of size *n* from a large population evolving according to the Wright-Fisher model (Ewens, 2004). The ancestry of this sample is described by its genealogy (Kingman, 1982a,b), and *K* represents the number of mutations that occurred along the sample’s genealogy. While neither the genealogy nor the specific times and locations of these mutations are directly observed, *K* can be identified as the number of polymorphic (segregating) sites under the infinite-sites model. *K* can be interpreted as: (1) the sum of mutations across different branches of the genealogy; (2) the sum of mutations classified by their size, i.e., based on the number of mutants in the sample (Fu, 1995); or (3) the sum of mutations that occurred across the ancestral sequences in different states of the coalescent process. It is this latter partition of *K* that we will explore to derive its asymptotic distribution. Figure 1 illustrates this partition in the genealogy of a sample of 5 sequences.

**Fig. 1.**
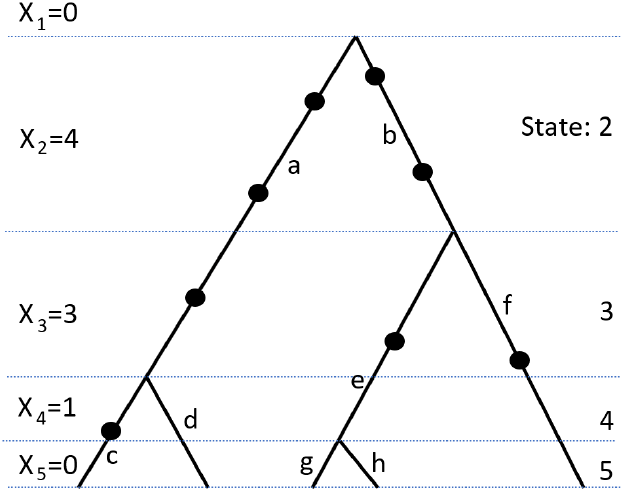
A genealogy of a sample of size 5 with mutations marked by black dots. Mutations can be classified in several ways: (1) by the state of coalescent process at which they occur, (2) by the branch on which they occur, or (3) by the number of mutants (mutation size) they result in. The first classification leads to *X*_*i*_ 0, 4, 3, 1 and 0 respectively for *i* 1, …, 5, the second leads to *a* 3, *b* 2, *c* 1, *d* 0, *e* 1, *f* 1, *g* 0 and *h* 0, and third leads to mutations of size 1 is 2, of size 2 is 4 and of size 3 is 2.

Looking backward in time, the number of ancestors of the sample decreases from *n* to *n*−1, then *n*−2, and eventually to one, which is the most recent common ancestor of the sample. Let *t*_*k*_ denote the coalescent time (i.e., the waiting time) for *k* ancestors to coalesce into *k*−1 (scaled such that one unit corresponds to 4*N* generations). According to coalescent theory (Kingman, 1982a,b), *t*_*k*_ is exponentially distributed with parameter *k* (*k*−1). Although Kingman’s coalescent assumes that the sample size *n* is much smaller than the population size under the Wright-Fisher model, it is well known that the results are robust across several population genetics models and are exact under the Moran model, without any restriction on sample size (Ewens, 2004). Therefore, we assume throughout this paper that exponentially distributed waiting times apply to all sample sizes. With the notation 𝔼 (or 𝔼 []) representing the expectation, we are now ready to prove the central limit theorem for *K*.

### Theorem.

*Suppose K is the number of mutations in the genealogy of a sample of size n from a large population that evolves according to the Wright-Fisher model with a constant effective size N. Then K 𝒩* ∼ *(a*_*n*_*θ, a*_*n*_*θ + b*_*n*_*θ*^2^) *as n* ⟶ ∞, *where θ* 4*Nµ with µ as the mutation rate per sequence per generation, and a*_*n*_ *and b*_*n*_ *defined by (3) and (4), respectively*.

**Proof**. The genealogy of a sample of size *n* can be partitioned into *n−* 1 segments based on the states (i.e., the number of ancestors) of the coalescent process (see Fig. 1). Let *X*_*k*_ denote the number of mutations that occur during state *k* (*2*⩽ *k* ⩽ *n*), and for convenience, we define *X*_1_ = 0. Then

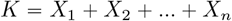

With the convention that the number of mutations in a lineage (branch segment) of given its length *l* (scaled so that one unit corresponds to 4N generations) is a Poisson variable with parameter *lθ*, it follows for *k ⩾* 2 that *X*_*k*_ ∼ *Pois*(*kθt*_*k*_) where *t*_*k*_ ∼ *Exp*(*k(k* − 1)), which is the kth coalescent time. It follows that *X*_*k*_ ∼ *Geom*p*p*_*k*_q where 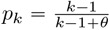, which led to that for 2 ⩽ *k*

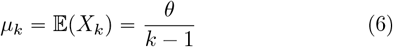

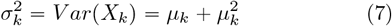

and that 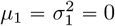. Since *X*_*k*_ are mutually independent for different *k*, the mean and variance of *K* are respectively the sum of those for *X*_*i*_ (*i* = 1, …, *n*). That is

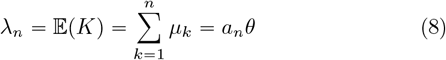

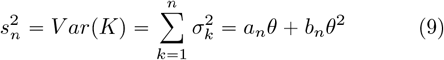

where *a*_*n*_ and *b*_*n*_ are given by (3) and (4) respectively. Furthermore *a*_*n*_ is non-convergent with lim_*n*→ ∞_ [*a*_*n*_ − ln (*n* − 1] = *γ* where *γ* is Euler constant (Abramowitz and Stegun, 1972), while lim_*n*→ ∞_ *b*_*n*_ = *π*^2^ 6.

The Central Limit Theory (CLT) has several versions (Feller, 1968). According to the Lindegerg Central Limit Theorem (Lindeberg (1922), also see page 369 in Billingsley (1986) and page 254 in Feller (1968)), a sufficient condition for

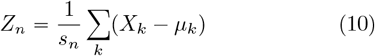

to follow asymptotically *N* p0, 1q is that for every *ϵ* > 0

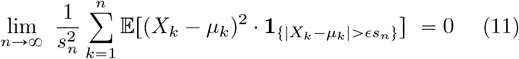

where **1**_{…}_ is the indicator variable, which takes value 1 when the specified condition is satisfied and 0 otherwise.

Since 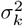 is monotonically decreasing with *k* and *s*_*n*_ is non-convergent, it follows that for any *ϵ >* 0, *ϵs*_*n*_ will eventually be larger than all *u*_*k*_, (*k* = 1,…). Consequently for a sufficiently large *n*, the event {|*X*_*k*_ − *µ* |*> ϵs*_*n*_ = {*X*_*k*_ − *µ*_*k*_ > *ϵs*_*n*_, which is a subset of {*X*_*k*_ ⩾ [*ϵs*_*n*_]} where [*x*] stands for the smallest integer greater or equal to *x*. Let *m* = [*ϵs*_*n*_] and 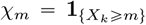, it follows that for a sufficiently large *n*

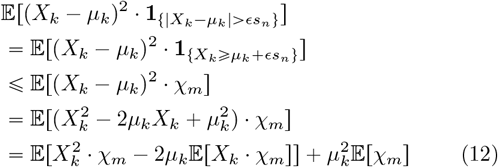

Since *X*_*k*_ is a geometric random variable with 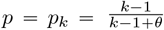, it follows that

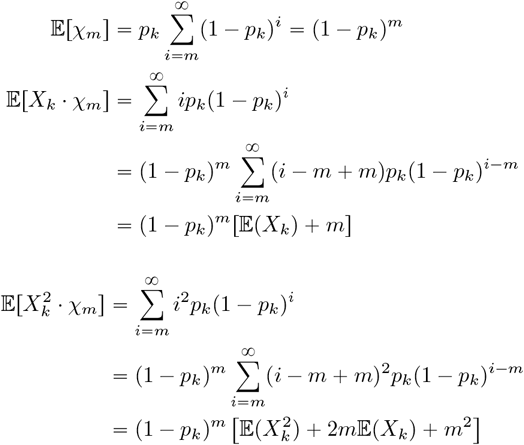

Substituting these results for corresponding terms in (12) together with Eq.(6) results in

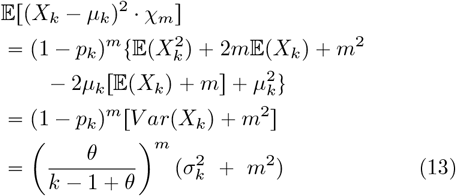

Since 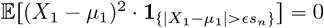, it follows that

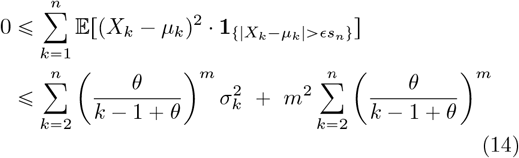

Since 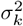 is monotonically decreasing, the first summation is smaller with 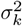 replaced by 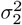, therefore convergent as long as *m* > 1. For the second summation, one can decouple the dependence of the number of summands and *m* by noticing

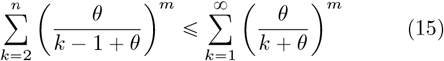

Denoting the right hand side of above inequality as *ν*(*m, θ*) and since for every integer *n*^′^ > 1

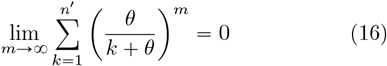

therefore lim_*n*→∞_ *ν* (*m, θ*) lim_*m*→∞_ *ν* (*m, θ*) = 0, which means that

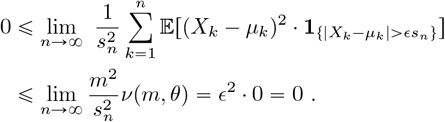

Therefore the Lindeberg condition (11) is satisfied, which implies that *Z*_*n*_ *∼* 𝒩(0, 1) as *n* → ∞. Since

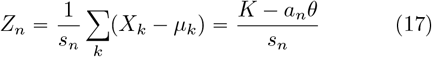

Therefore 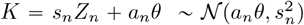 for sufficiently large *n*. ‖

We now consider a scenario in which the effective population size remains constant within each state of the coalescent process of a sample, while the effective population sizes across different states can be different but are mutually independent. This model will be referred to as the *constant-in-state model*. The constant-in-state model serves as a straightforward extension of the Wright-Fisher model with a constant effective population size. It offers mathematical simplicity while allowing considerable flexibility in modeling population size dynamics. Consequently, this model has been employed in inferring historical population size changes (Pybus et al., 2000; Liu and Fu, 2015).

Let *N*_*k*_ be the effective population size at state *k* of the coalescent and *θ*_*k*_ = *N*_*k*_*µ*. Assume that *N*_*k*_ are uniformly bounded such that there exist finite constants *θ*_*min*_ and *θ*_*max*_ such that for every *k* > 1

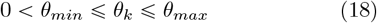

Furthermore assume that *X*_*i*_ are independent for different *i*. Thus under the constant-in-state model, Eqs.(6 - 9) naturally extend to

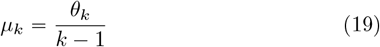

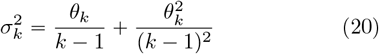

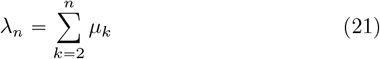

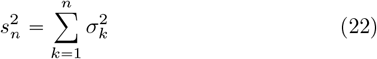

With these modifications, it follows that Eqs. (13) - (15) continue to hold by substituting *θ*_*k*_ for *θ*. Furthermore the limit in Eq.(16) is also true regardless of *θ* being constant or not, as long as its replacement *θ*_*k*_ are uniformly bounded, which also guarantees that *s*_*n*_ is non-convergent so that the first summation in (14) divided by *s*_*n*_ will be zero as *n* → ∞. As *K* = *s*_*n*_*Z*_*n*_ +*λ*_*n*_, it follows that

### Corollary

*For a large population evolving according to the constant-in-state model such that the effective size N*_*k*_ *for state k* (*k* = 2, …) *are bounded uniformly*, 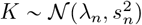 *as n* → ∞, *where λ*_*n*_ *and* 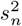 *are defined by (21) and (22), respectively*.

It should be emphasized that the proofs of the Theorem and Corollary establish only the asymptotic normality for *Z*_*n*_. Since lim_*n*→ ∞_ = ∞ and lim_*n*→ ∞_ ∞, it is technically incorrect to assert that the limiting distribution of *K* is normal. Therefore the asymptotic normality of *K* should be interpreted as approximate normal for large sample when the normality of *Z*_*n*_ is justified.

## The Skewness and Kurtosis of *K*

Since *K* and *θ*_*W*_ are prevalent in both theoretical and data analysis contexts, complete characterizations of these statistics are highly desirable. This necessitates the examination of higher moments, particularly skewness and kurtosis, with the former of which measuring symmetry while the later tailedness (Westfall, 2014). It is important to note that it is sufficient to derive the skewness and kurtosis of *K*, as these two quantities are invariant under positive scaling.

Under the constant-in-state model, *X*_*i*_ follows geometric distribution with *p* = *p*_*i*_ = p*i* = (*i* − 1) /(*i* − 1 + *θ*_*i*_) and is independent of *X*_*k*_ (*k* ≠ *i*). The moment generating function of *K* is thus the product of those of *X*_*i*_ (*i* = 2, …, *n*)

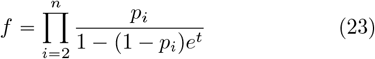

Consequently

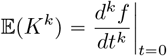

It is thus straightforward to derive the power moments although the process becomes tedious and lengthy with increasing *k*. For conciseness, define a zeta function

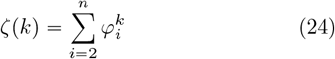

where *φ*_*i*_ = *θ*_*i*_/(*i* − 1). For a population of constant size such that *θ*_*i*_ = *θ*, it follows that

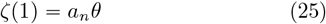

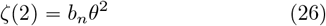

where *a*_*n*_ and *b*_*n*_ are given by (3) and (4) respectively. Thus *ζ (*1) is linear and *ζ (*2) is quadratic with regard to *θ*. If the power order of a polynomial of *θ*_*i*_’s is defined as the maximum number of *θ*’s in a product term, then *ζ (k)* is of power order *k*. In fact, every term of *ζ (k)* is of power order *k*. Consequently *ζ (i*_1_) *ζ (i*_2_)…*ζ (i*_*k*_) is of power order *i*_1_ +…+ *i*_*k*_.

The first four power moments of *K* are (from Eqs (35)-(38) in Appendix)

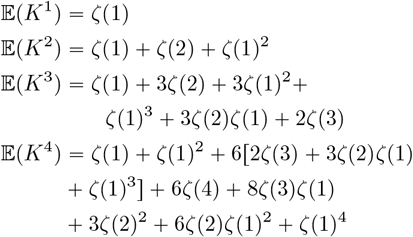

In general it appears that 𝔼(*K*^*k*^) is of power order *k*. The central moments of *K* can be obtained through

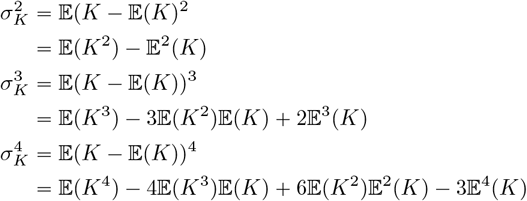

Simplification after substituting 𝔼(*K*^*i*^) by their expressions leads to

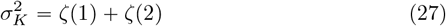

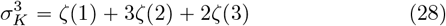

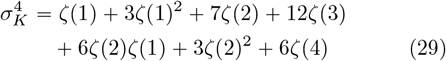

It is obvious that when the population size is constant, 𝔼(*K*) and 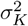 will be the same as (1) and (2), respectively. The above results also suggest that the *k*-th central moment is of power order *k*. With 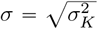, the skewness *γ(K)* and kurtosis *κ(K)*, which are the 3rd and 4th standardized moments respectively, become

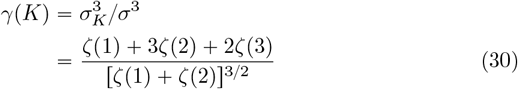

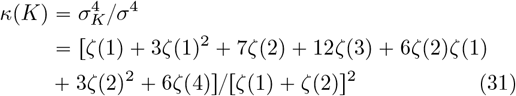

It is clear that *γ*(*K*) is always positive, which indicates a longer tail for large values of *K*. Since both *ζ*(2) and *ζ*(3) are convergent with *n*, thus for a sufficiently large sample size

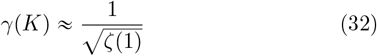

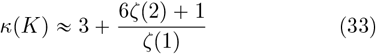

which will approach 0 and 3, respectively, as *n*→ ∞, which are the skewness and kurtosis for a normal distribution. These results thus agree with the now-established asymptotic normality of *K*. The convergent speed for *γ*(*K*) is at the order 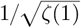 while that for *κ (K)* is at 1/ *ζ (*1). Intuitively the speed of approaching normality for *K* should be at most as faster as the convergent speed of standardized moments. Therefore, the speed of approaching normality for *K* should be at most as fast as 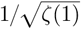. Under the model of constant size, *ζ*(1) = *a*_*n*_*θ* ≈ In(*n*)*θ*, the convergent speed for *γ (K)* is thus at the order 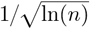, and that for *κ*(*K*) is at 1/ ln(*n*).

Fig. 2 presents the skewness and kurtosis of *K* in the case of a constant effective population size for several values of *θ* and a wide range of sample sizes. The most apparent pattern is that the convergences to the asymptotic skewness and kurtosis are relatively slow after initial rapid decreases, which agree with the earlier speed analysis. It is also obvious that kurtosis converges faster than skewness, agreeing with convergent speed 1/ ln *n* and 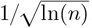, respectively, for kurtosis and skewness. Another noteworthy observation is that the trends for both skewness and kurtosis are not linear with regard to *θ*, with *θ* 1 yielding the smallest skewness and kurtosis for while range of sample sizes. Why *θ* 1 posses such characteristics is not obvious from either (30)-(31) or (32)-(33). However, since the dominant factor is 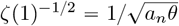 in *γ(K)* and is *ζ (*1) ^−1^ = 1 /(*a*_*n*_*θ)*, eventually both *γ K* and *κ K* will be smaller for larger *θ*.

**Fig. 2.**
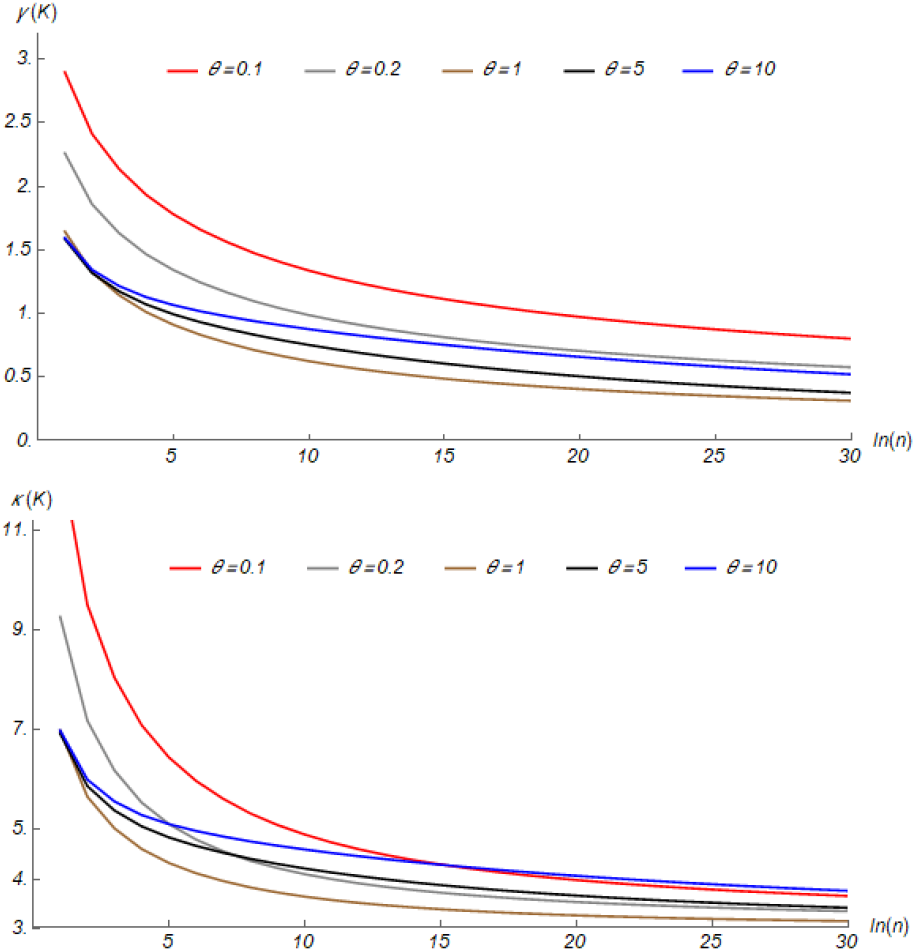
Skewness *γ (K)* (top panel) and Kurtosis *κ(K)* (low panel) of *K* with different values of *θ* and sample sizes (in natural logarithmic scale).

Fig 3. presents visual comparison of the distribution of *K* and normal distribution under the assumption of constant effective population size. For each *θ* examined, the improvement of normality from sample size 50 to 500 is visually apparent, but the changes from sample size 500 to 5000, is more subtle. In the case *θ =* 1, *γ (K)* for sample size 50, 500 and 5000 are, respectively, 0.78,0.57 and 0.47. The successive percentages of reduction are 27% and 18% respectively, which explains why the improvement of symmetry from 500 to 5000 is not profound. In comparison, the excess of kurtosis (i.e., *κ(K)* − 3) are 0.99, 0.55 and 0.36, respectively, for the three sample sizes, with successive percentages of reduction as 44% and 35%, which are nearly twice as large as those for skewness. As a result, the non-normal kurtosis of the distribution of *K* appears not as serious as its non-symmetry. Similar patterns are also observed in the case *θ =* 3. For the three sample sizes, *γ (K)* are, respectively, 0.86,0.62 and 0.50, while *κ (K)* are, respectively, 1.31, 0.84 and 0.59. However, one should keep in mind that skewness around 0.5 and excess of kurtosis less than 1 are generally considered as very close to normality (Hair et al., 2010)).

**Fig. 3.**
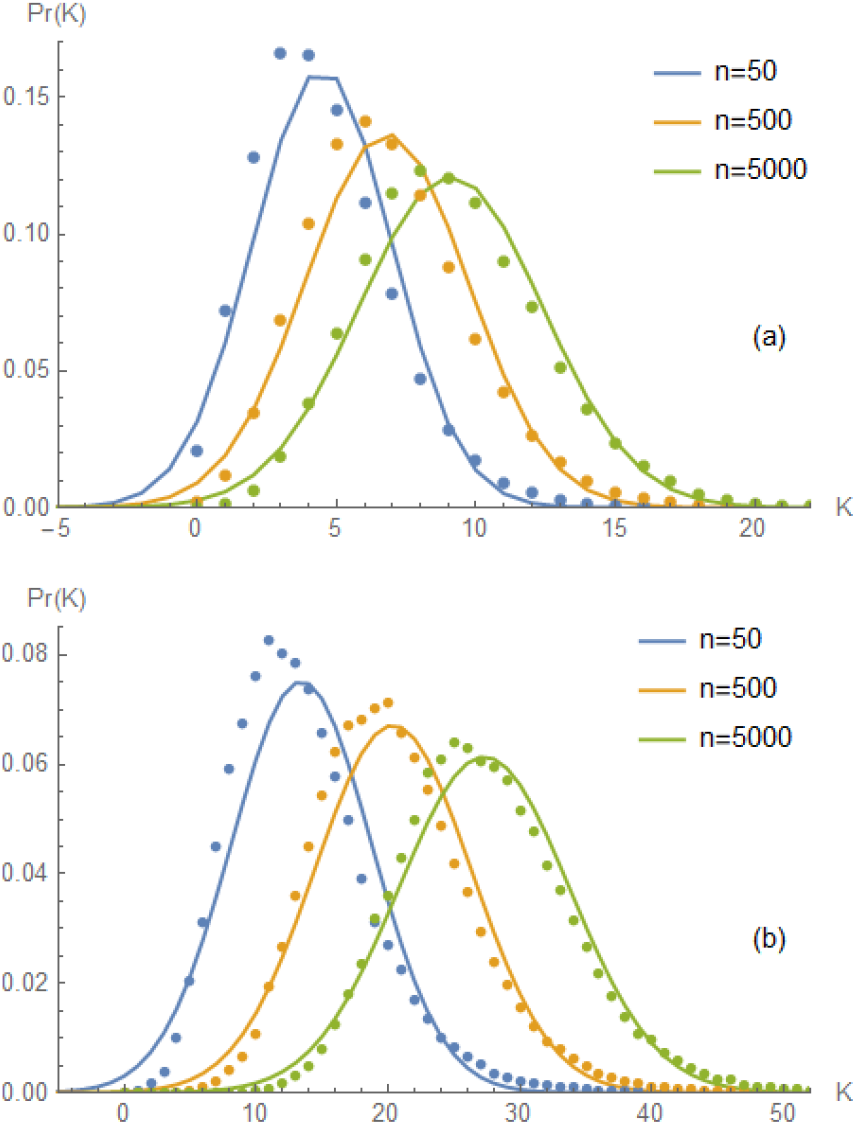
Distribution of *K* for three different sample sizes under the assumption of constant effective population size. Panel (a) *θ =* 1 and (b) *θ=* 3. Dots represent the distributions based on coalescent simulations with 50, 000 replicates (for each case) and smooth curves are the (discretized) normal distributions *𝒩* (*a*_*n*_*θ, a*_*n*_*θ* + *b*_*n*_*θ*^2^)

## Potential applications of asymptotic normality

Although the probability distribution of *K* for a finite sample, assuming a constant population size, is known (Eq, 9.5 in Tavaré (1984)),using Tavaré’s formula can be inconvenient even for modest sample sizes due to the accumulation of errors in the summation of many large summands with alternating signs. The asymptotic normality of *K* and *θ*_*W*_ suggests that normal distributions may be used to approximate its distribution. While the exploration of skewness and kurtosis in the previous section indicates that the convergence to normality is relatively slow, it remains meaningful to explore when normality is sufficiently accurate. The answer to this question is unlikely to be universal for all applications. Therefore, we investigate this issue from the perspective of whether normality can be rejected by applying Shapiro-Wilk test (Shapiro and Wilk, 1965), which is the most powerful existing test for such purpose (Razali and Wah, 2011).

There are two ways a sample of *K* can be obtained in practice. One is to take multiple samples of the same size from the same locus, each yielding an observation of *K*. This does not generate a proper sample of *K* for the purpose of evaluation, because different samples are in fact not independent, leading to correlated *K*’s which will complicate the analysis. Alternatively, one can examine the *K*’s for the same sample across different loci that are sufficiently apart such that the assumption of independence among loci are appropriate. This mimics the reality better since there have been multiple studies in which DNA polymorphism in a sample from a population were determined in large genomic regions or even the entire genome. Therefore, we decided to simulate samples of the same size from independent loci and examined how sample size and the number of loci affected the normality assessment.

Table 1 shows the rates of rejecting normality for some combinations of *θ*, sample size *n* and the number of loci, where each locus provides one observed value *K*. Interpretation of rejection rate should take into consideration that it generally increases with the number of loci while holding sample size constant, and decreases with sample size when holding the number of loci constant. This is because for a given sample size, more independent observations of *K* will lead to higher power of rejection of normality since normality is approximate at best, while for the same number of observations of *K*, larger sample sizes will improve the normality approximation, and thus lead to reduced rejection rate. This is what Fig. 3 suggests and what Table 1 reveals. For example, when *θ* = 10 and *n* = 100, three numbers of loci, 50,100 and 200, lead to rejection rates (at 1% significance level) 30%, 76% and 96%, respectively. On the other hand, when *n* = 1000, the same numbers of loci lead to reject rates 24%,59% and 88%, respectively. The high rejection rate for *θ* = 0.1 with 50 loci also agrees with Fig.2. which shows severe skewness and kurtosis for *θ <* 0.5. Overall, a sample of 200 or more *K*’s will have quite high rejection rate regardless of sample size and *θ*.

**Table 1.** Rates of rejection of Normality by Shapiro-Wilk test for simulated samples.

| $\theta$ | $L^a$ | 1%<sup>b</sup> | 5%<sup>c</sup> | $\theta$ | $L^a$ | 1%<sup>b</sup> | 5%<sup>c</sup> |
| --- | --- | --- | --- | --- | --- | --- | --- |
| n=100 |  |  |  | n=1000 |  |  |  |
| 0.1 | 50 | 100 | 100 | 0.1 | 50 | 100 | 100 |
| 0.5 | 50 | 80 | 99 | 0.5 | 50 | 44 | 76 |
| 1.0 | 50 | 40 | 66 | 1.0 | 50 | 11 | 41 |
|  | 100 | 82 | 97 |  | 100 | 42 | 76 |
| 5.0 | 50 | 24 | 41 | 5.0 | 50 | 21 | 35 |
|  | 100 | 54 | 78 |  | 100 | 40 | 54 |
|  | 200 | 88 | 99 |  | 200 | 76 | 90 |
| 10 | 50 | 30 | 60 | 10 | 50 | 24 | 40 |
|  | 100 | 76 | 88 |  | 100 | 59 | 77 |
|  | 200 | 96 | 97 |  | 200 | 88 | 97 |
| n=500 |  |  |  | n=5000 |  |  |  |
| 0.5 | 50 | 49 | 83 | 0.5 | 50 | 23 | 65 |
|  | 100 | 96 | 100 |  | 100 | 81 | 99 |
| 1.0 | 50 | 22 | 46 | 1.0 | 50 | 19 | 34 |
|  | 100 | 61 | 88 |  | 100 | 33 | 65 |
|  | 200 | 97 | 100 |  | 200 | 88 | 99 |
| 5.0 | 50 | 15 | 34 | 5.0 | 50 | 14 | 26 |
|  | 100 | 44 | 63 |  | 100 | 30 | 49 |
|  | 200 | 78 | 92 |  | 200 | 55 | 77 |
| 10 | 50 | 28 | 46 | 10 | 50 | 25 | 41 |
|  | 100 | 54 | 77 |  | 100 | 53 | 64 |
|  | 200 | 92 | 98 |  | 200 | 85 | 96 |

<sup>a</sup>: The number of loci, i.e., the number of $K$ sampled; <sup>b</sup>: rejection rate ( $\times 100$ ) at 1% significance level and <sup>c</sup>: rejection rate ( $\times 100$ ) at 5% significance level. For each combination of $n, \theta$ and $L$ , results are based on 100 replicates of coalescent simulation.

An reasonable procedure in data analysis may be as follow: When a modest sample of *K* is collected, Shapiro-Wilk test will be applied and when normality fails to be rejected, one then proceeds to use a normal distribution for the sample of *K*^1^*s* for subsequent analysis. Results in Table 1 thus suggest there will be ample opportunities for asymptotic normality of *K* to play a role, particularly when the number of loci is less than 200 and the finite sample exact distribution is not available or difficult to use.

## Discussion

It is now established that the Central Limit Theorem holds for *K* in a large population evolving according to the Wright-Fisher model with a constant effective size, as well as under the uniformly bounded constant-in-state model.However, the skewness and kurtosis of *K* indicate that the speed of convergence to asymptotic normality is relative slow after an initial rapid decrease, which limits the practical utility of asymptotic normality. This outcome is not too surprising given that *V ar θ*_*W*_ approaches the minimal variance of estimators of *θ* rather slowly (Fu and Li, 1993a). In the most favorable situation, the mean of *n* independent and identically distributed random variables approaches normality at the speed 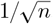, while *K* ‘s speed to normality is at best 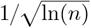 (a formal analysis may be achievable using Berry-Esseen theorem, page 544 in Feller (1971)).

It should be pointed out that our intuitive analysis on whether asymptotic normality for *K* can play a role in practice is based on very stringent criteria— passing the normality test by Shapiro and Wilk test. In statistical analysis, often the rule of thumb (Hair et al., 2010) for accepting normality is that skewness is within -2 and +2, and excess of kurtosis is within -7 and +7. If such liberal standard is adopted, then normality for almost all cases considered in Table 1 will be acceptable. As a whole, it is probably reasonable to conclude that although the convergent speed to asymptotic normality for *K* is relatively slow, its distribution for a reasonable sample size is already quite close to normality.

The higher moments, particularly skewness and kurtosis, of *K* may have practical utility. On one hand, they can be employed to assess the characteristics of polymorphism in DNA samples against theoretical expectations. On the other hand, they may be useful for exploring summary statistics that exhibit desirable properties, similar to linear functions of the number of mutations of various sizes (e.g., Fu (2022)). Finally, higher moments provide opportunities to model the probability distributions of *K* under various assumptions, which might offer computational advantages over exact distributions (if they can be derived).

## Appendix

### Derivation of 𝔼 (*K*^3^) and 𝔼 (*K*^4^)

The moment generating function of *K* under the assumption that *X*_*i*_’s are mutually independent for differnt *i* can be written as

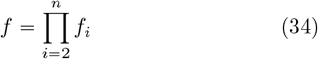

where *f*_*i*_ = *p*_*i*_*q*^−1^ with *p*_*i*_ = (*i* − 1)/(*i* − 1 + *θ*_*i*_) and *q*_*i*_ = 1 − (1 − *p*_*i*_)*e*^*t*^. Define 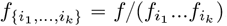, which is *f* with factors 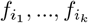 removed. With the convention that *x*^(*k*)^ represents the *k*-th derivative of *x*, one can obtain derivatives of *f* sequentially. The first four derivatives are

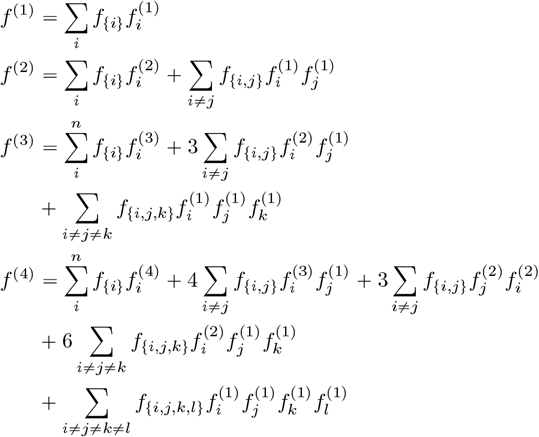

While for a given *i*

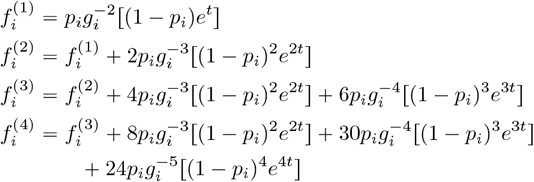

where *g*_*i*_ =1 − (1 − *p*_*i*_)*e*^*t*^. Let 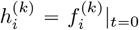. It follows that

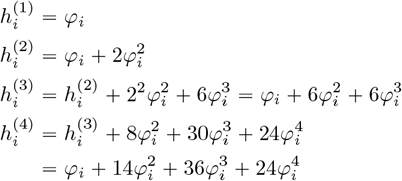

In addition to the relationship (25) and (26), it is straight-forward to verify that

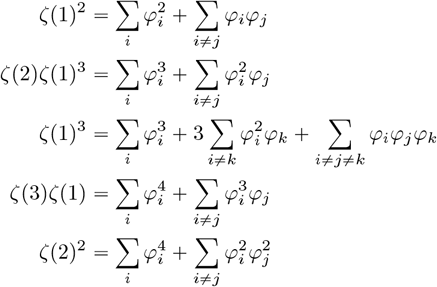

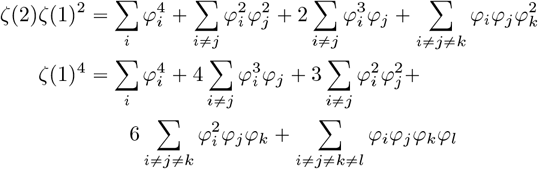

Since 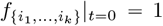, thus *f* ^(*k*)^|_*t*=0_ is ready to be obtained for *k* ⩽ 4. It follows that

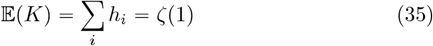

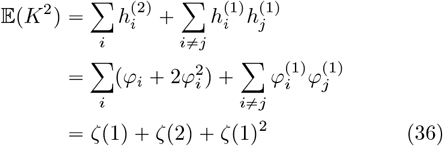

which are well known. It follows similarly that

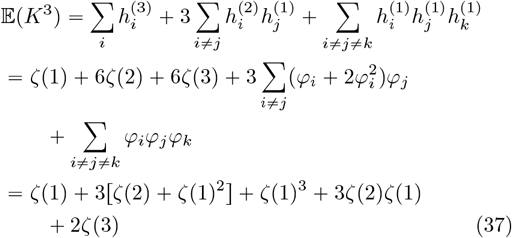

while

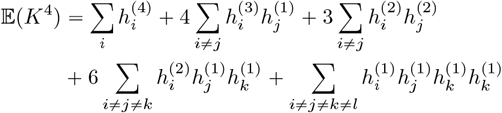

with

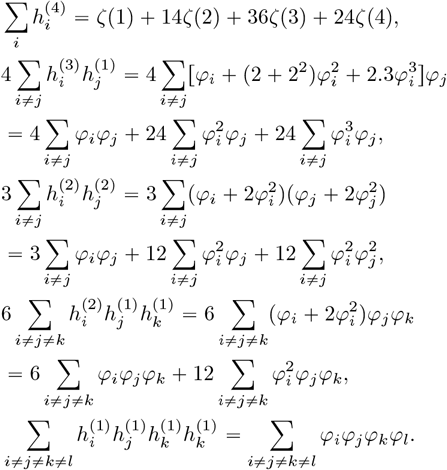

Grouping terms according to their power order will leads to concise expression. It is easy to see that the first power order term is *ζ*(1), the second order power term is *ζ*(1)^2^, the third power order term is

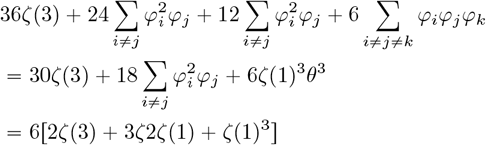

and the fourth power order term is

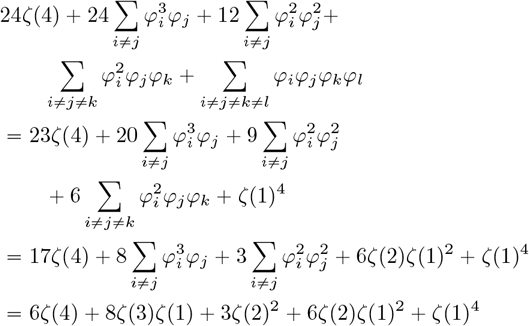

Therefore

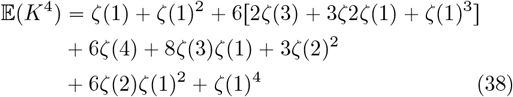

